# Blastocoel fluid RNA predicts pregnancy outcome in assisted reproduction

**DOI:** 10.1101/2025.03.09.641486

**Authors:** Caroline Frisendahl, Kristina Gemzell-Danielsson, Karthik Nair, Forrest Koch, Judith Menezes, Mona Sheikhi, Karin Dahlgren, Wesam Alnahal, Karin Lindgren, Therese Bohlin, Jan I. Olofsson, Virpi Töhönen, Fatemeh Vafaee, Parameswaran Grace Lalitkumar, Nageswara Rao Boggavarapu, Omid R. Faridani

## Abstract

Nearly one in eight couples are affected by infertility, with many relying on assisted reproductive technologies (ART) to conceive. However, selecting the highest-quality embryo in ART remains a major challenge, as current assessment methods are often subjective, or invasive, and lack precision. Here, we introduce a novel strategy that analyses embryo-derived polyadenylated RNA in blastocoel fluid to more accurately predict pregnancy outcomes. Elevated RNA levels were strongly associated with implantation failure, particularly in embryos from women over the age of 34. Our predictive model developed using our sample cohort, incorporating both RNA and maternal age, demonstrated exceptional performance, achieving 76% accuracy in the training set and 73% in independent validation in predicting implantation outcome— highlighting a promising advancement in embryo selection and ART success.

## Main

Despite multiple ART cycles, more than one-third of couples do not achieve a live birth, highlighting the limitations of current assisted reproductive technologies^1^. A major challenge in ART is identifying an embryo with the greatest potential to result in a healthy live birth. While morphological grading has been used for embryo quality assessment for over the past four decades, it is assessor-dependent and often fail to reliably predict treatment outcomes^2^. The application of artificial intelligence (AI) algorithms leveraging morphokinetic and morphological data derived from time-lapse imaging has shown promise in enhancing the objectivity of embryo assessment. Despite these exploratory advancements, prospective studies have yet to demonstrate that these AI-driven methods outperform conventional morphological scoring in clinical outcomes^3,4^. Preimplantation genetic testing for aneuploidy (PGT-A) may complement embryo assessment, but it is costly, time-consuming, requires specialised expertise, and the long-term effects of the invasive trophectoderm biopsy remains unclear^5,6^. PGT-A does not improve live birth or reduce miscarriage rates for most IVF patients, especially younger women or those without recurrent pregnancy loss or advanced maternal age^7^. Its benefits appear limited to specific subgroups, such as women aged 35 and older^7^.

Embryo spent culture media (ESM), that contains cell-free molecules including RNA, DNA, metabolites, and proteins, has been investigated as a source of biomarkers to assess embryo quality objectively and non-invasively^8–11^. Despite their promise, metabolite-, protein-, and microRNA-based biomarkers identified in ESM have demonstrated limited clinical translation and inconsistent results. This is likely attributable to technical challenges, specifically the difficulty of analysing target molecules within the very small volume of ESM obtained from individual embryos^8,11–14^.

Additionally, published microRNA studies that utilised pooled ESM samples are limited in their ability to predict pregnancy outcomes for individual embryos^8,12,14^. Cell-free DNA in ESM has been investigated as a non-invasive method for PGT-A, showing comparable aneuploidy detection rates; however, it remains unclear whether incorporating this method will improve pregnancy outcomes^9^. A recent study suggested that extracellular mRNA in ESM is embryo-related and undergoes dynamic changes during development^10^. Blastocoel fluid (BF) is a promising source for embryo quality markers. Its unique composition, reflecting the blastocyst’s internal environment, offers a valuable opportunity to discover novel biomarkers, potentially leading to more accurate embryo assessment^15,16^. Our study is the first to investigate extracellular polyadenylated RNAs in both ESM and BF as objective and non-invasive biomarkers of embryo quality in ART.

We developed an innovative RNA library preparation protocol for sequencing of microlitre-scale fluids enabling individual analysis of low volume and low RNA content samples from cultured pre-implantation embryos (Fig. 1a). In this prospective observational multicentre study, ESM samples were collected from embryos either at cleavage stage (day 3) or blastocyst stage (day 5/6) immediately after fresh embryo transfer or vitrification for later cryopreserved transfers. BF samples were obtained on culture day 5 or 6 by laser collapsing blastocysts into fresh medium before vitrification and later frozen transfer (Fig. 1a). The quality of all the embryos in the study was assessed morphologically by an embryologist before transfer or vitrification. A pregnancy test based on urinary human chorionic gonadotropin (hCG) levels was used to determine implantation outcome following embryo transfer. We also collected data on live-birth and negative pregnancy outcomes including biochemical pregnancy and spontaneous miscarriages. All ESM and BF samples were subjected to RNA sequencing. We analysed the RNA content from ESM samples of 105 cleavage-stage and 230 blastocyst-stage embryos following fresh or frozen-thaw transfers, as well as from BF samples of 72 earlier cryopreserved blastocyst-stage embryos that upon thawing underwent transfer. Clinical characteristics of the included samples can be found in Extended Table. 1.

**Fig 1.**
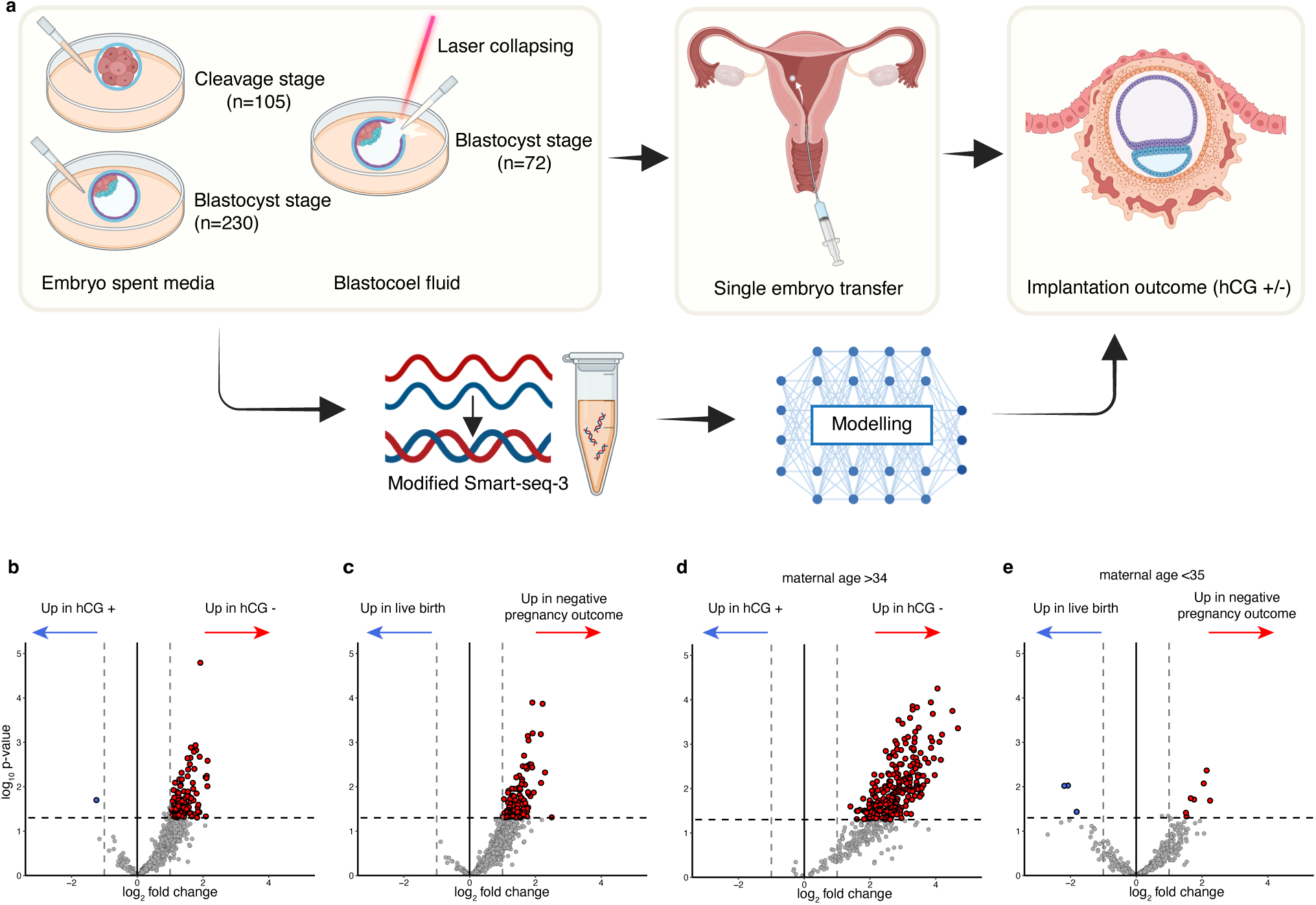
Detection of polyadenylated RNA associated with pregnancy outcomes in blastocoel fluid. **a**, Flow-chart of the study design. **b-e**, Volcano plots demonstrating differences in blastocoel fluid RNA between: hCG negative and hCG positive blastocysts (n=72) (**b**); negative pregnancy outcome and live-birth blastocysts (n=68) (**c**); hCG negative and hCG positive blastocyst from women > 34 years of age (n=29) (**d**); and negative pregnancy outcome and live-birth blastocysts from women <35 years of age (n=43) (**e**). Red and blue dots represent significantly upregulated and downregulated genes respectively detected using DESeq2 (p-value <0.05 and a log2 fold change ≤-1 or ≥1). Grey dots represent unsignificant genes. Abbreviations: hCG=human chorionic gonadotropin. Flow-chart created with BioRender. Alkasalias, T. (2025) https://BioRender.com/sb7koyg.

RNA quality assessed by the average Transcript Integrity Number (TIN), was higher in BF samples compared to ESM samples (Extended Fig. 1a). TIN value reflects RNA integrity by measuring how uniformly sequencing reads cover each transcript^17^. A substantial number of polyadenylated RNAs were identified in ESM and BF samples collected from embryos of varying developmental stages (Extended Fig. 1b). The majority of these polyadenylated RNAs were common in ESM samples from both cleavage-stage and blastocyst-stage embryos as well as in BF samples (Extended Fig. 1b).

To investigate whether extracellular polyadenylated RNA content correlates with pregnancy outcomes, we conducted differential gene expression analysis. We compared the RNA content of samples associated with embryos that resulted in no implantation (hCG-negative) or implantation (hCG-positive), as well as embryos that resulted in negative pregnancy outcomes (including miscarriages and biochemical pregnancies) with those that led to live births.

In ESM samples collected from cleavage- and blastocyst-stage embryos, we identified few differentially abundant RNAs when comparing hCG negative with hCG positive embryos (Extended Fig. 2a-b). Similar results were obtained when comparing the RNA content of ESM samples from cleavage- and blastocyst-stage embryos with negative pregnancy outcomes to those resulting in live births (Extended Fig. 2c-d). Further subgroup analysis of the ESM samples, based on different blastocyst developmental stages, revealed minimal differences in RNA content between hCG positive and hCG negative embryos as well as between negative pregnancy outcomes and live birth embryos (Extended Fig. 2e-l). Overall, extracellular RNA content in ESM did not strongly correlate with pregnancy outcomes.

In BF samples, however, a significant increase in abundancy of polyadenylated RNA was identified in hCG negative compared to hCG positive blastocyst embryos (Fig.1b). Similarly, significantly more polyadenylated RNA were detected in BF samples collected from embryos that did not result in a live birth (Fig. 1c). Collectively, the results demonstrate that RNA content in BF samples has a stronger association with pregnancy outcomes compared to that in ESM. Next, we investigated the influence of maternal age on the RNA content of BF samples, given that advanced maternal age is a primary risk factor for embryonic aneuploidy and subsequent first-trimester miscarriages^18,19^. Interestingly, embryos from women older than 34 years showed higher RNA levels in hCG negative samples, a trend not seen in younger women (Fig. 1d-e). A comparable but weaker age-related trend was observed in embryos that did not result in a live birth, suggesting that maternal age may influence the RNA profile of BF samples, particularly in low quality embryos (Extended Fig. 3a-b).

To identify the extracellular RNA signature that predicts pregnancy outcome, we developed a Leave-One-Out Cross-Validation (LOOCV) machine learning model based on the polyadenylated RNA detected in BF samples. This strategy was used to allow for optimisation of the model despite the limited number of samples. The BF cohort was divided into a training and testing set (n=50) and an independent validation set (n=22). As maternal age influenced the RNA profile in BF, we included age as an additional feature in the prediction models tested. The performance of the best prediction model, top three prediction models and other tested models are presented in Fig 2a, Extended Fig. 4, and Extended Table 2 respectively. The most optimal prediction model used L1 regularised logistic regression, genes with RNA counts greater than 5 and maternal age as input features, with ‘scale’ as the preprocessing step (Fig. 2, Extended Table 2). Any RNA detected in water or medium-only control samples were excluded from the model. In total 225 genes and maternal age contributed with weightage in the model. The model predicted implantation outcomes with a sensitivity of 81%, specificity of 71%, an area under the curve (AUC) of 0.86, and an overall accuracy of 76% (Fig. 2a, Extended Table 2). Validation of the model in an independent set of BF samples achieved both a sensitivity and specificity of 73% in predicting implantation outcome (AUC 0.73 and accuracy of 73%) (Fig 2b).

**Fig 2.**
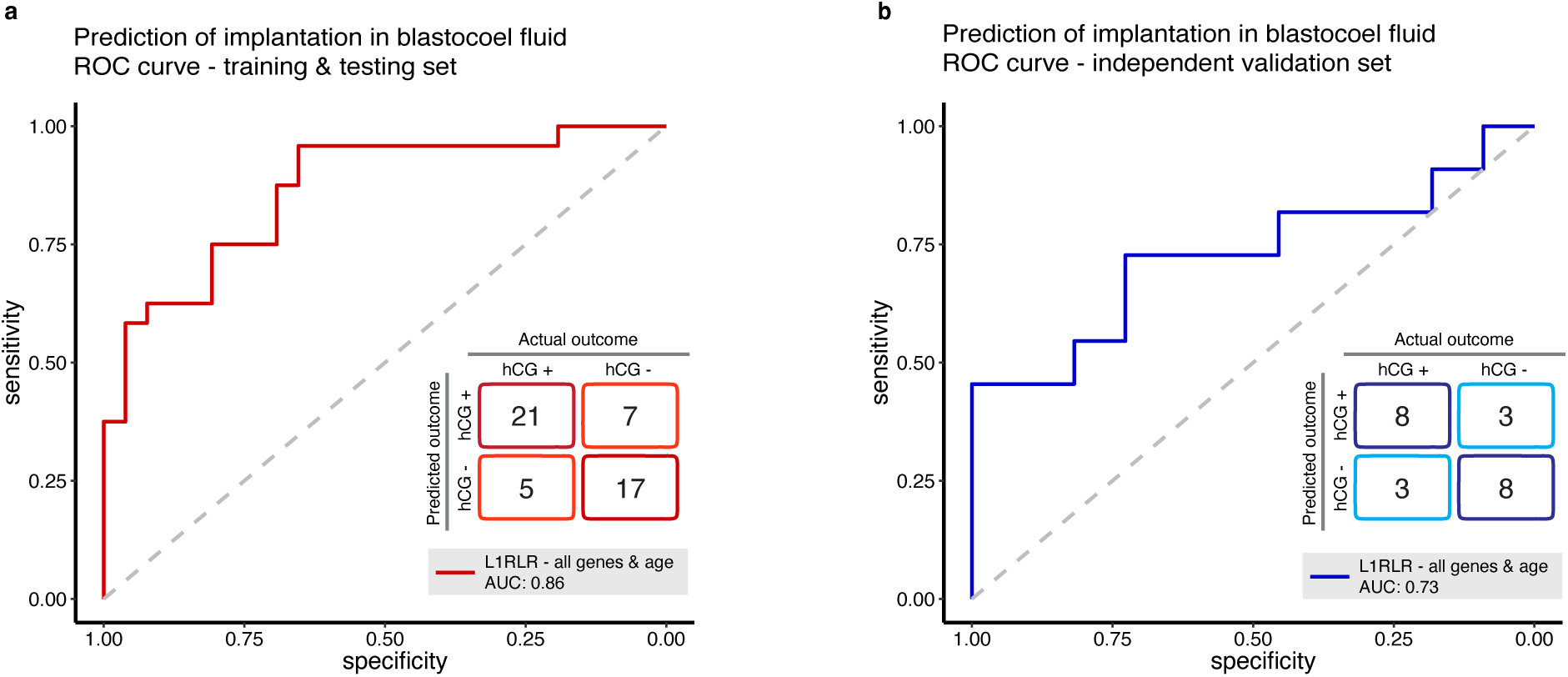
Prediction of embryo implantation based on blastocoel fluid RNA and maternal age. **a-b**, ROC curves and confusion matrices of the most accurate prediction model of embryo implantation based on blastocoel fluid RNA and maternal age in the training and testing set (n=50) (**a**) and in the independent validation set (n=22) (**b**). The prediction model utilises L1RLR as model class, all genes and maternal age as input features and pre-processing using scaling. L1RLR was performed using Glmnet. Implantation was defined as a positive urine hCG 14 days after embryo transfer. Abbreviations: hCG=human chorionic gonadotropin, LIRLR=L1 regularised logistic regression; AUC=area under the curve.

Our RNA-based model should be viewed as complementary to morphological assessment, as all transferred embryos in the study were already classified as high quality based on morphology. To compare the performance of our RNA-based prediction model to morphological assessment, we ran the same prediction model using morphological grades as input features. The morphology-based prediction model showed lower performance, with an accuracy of 56%, a sensitivity of 77%, and a specificity of 33% (AUC 0.56) (Extended Fig. 4d-e, Extended Table 2). Including maternal age as a feature to the morphology-based model led to only a minor improvement in AUC to 0.59 and specificity to 50%, while sensitivity decreased to 62%, resulting in the same accuracy of 56% (Extended Fig. 4d, f, Extended Table 2).

This study finds that embryos with negative pregnancy outcomes contained significantly higher levels of polyadenylated RNA in their BF samples. A predictive model based on RNA in BF and maternal age accurately classified implantation outcome based on hCG results and was successfully validated independently. The correlation of RNA content and pregnancy outcomes observed in BF samples was not detected in ESM samples. This may be attributed to lower RNA integrity in ESM, as indicated by the lower TIN values observed in these samples. In contrast, immediate BF collection after embryo collapse in fresh medium preserves RNA integrity, making it a more reliable source for implantation-stage biomarkers.

Compared to our study, a previous study by Kirkegaard et al. was unable to relaibly detect mRNA in ESM and BF samples due to technical limitations, highlighting the importance of sensitive methods for low-volume and low RNA content samples^20^. While another study detected RNA in ESM samples, co-culture of embryos in that study made it impossible to link findings to individual embryo outcomes^10^. A recent prospective study suggested that DNA absence in BF samples may correlate with an increased likelihood of live birth, but its small sample size and lack of proper controls limit its validity^15^. Another study suggested that aneuploid cells may be expelled into BF as a self-correcting mechanism^21^. This is consistent with our finding of elevated RNA levels in BF samples from embryos with negative pregnancy outcomes, particularly in women over 34 years of age.

Machine learning models based on morphological and morphokinetic parameters alone or in combination have gained increasing attention for embryo selection. However, most of the published models based on retrospective data from transferred blastocyst embryos show lower predictive performance than the RNA-based model in this study^3,22–24^,. For instance, KIDScore and IDAScore predictive models achieved AUCs of 0.66 and 0.67, respectively^22^. One larger study using a random forest model, as well as a pilot study that employed a convolutional neural network but without validation, reported AUCs of 0.74 and 0.77, respectively^23,24^. IDAScore was validated in a double blinded randomised controlled trial and found to be non-inferior to standard care^4^. Nonetheless, the overall clinical benefit of such models remains uncertain.

Laser-assisted blastocyst collapse is a commonly used technique in many ART laboratories to prevent ice crystal formation during vitrification, making it easy to integrate BF RNA analysis into embryo assessment for frozen embryo transfers^25^. This novel RNA-based approach is particularly well-suited for a segmented cycle strategy, where all embryos undergo vitrification and embryo transfer is deferred to a subsequent cycle^26^.

This study has several limitations. The small sample size limits the robustness of the predictive model, despite the use of a leave-one-out cross-validation strategy to reduce overfitting. There is also an inherent selection bias in the study design, since only embryos already deemed high-quality based on morphology were transferred and thus included in the study. In addition, the morphology data and RNA molecules are retrospectively correlated to implantation outcome. Therefore, larger prospective clinical trials are needed to confirm our findings and compare this prediction model to standard morphological assessments. Furthermore, the absence of reliable markers for endometrial receptivity restricts the model’s predictive power. Nonetheless, the balanced clinical characteristics between the groups help mitigate the risk of major confounding effects.

If confirmed in larger prospective clinical trials, this minimally invasive objective RNA-based technique could significantly improve ART by identifying high-quality embryos, potentially enhancing pregnancy success rates while reducing overall costs and burden.

## Material and Methods

### Ethics

The study was approved by the Research Ethics Committee at Karolinska Institutet. Original approval (Dnr 00-134) and amendments (Dnr 2009/1615-32; Dnr 2012/1413-32, Dnr 2016/1290-32 and Dnr 2018/567-32). Collection of the included material has been performed after written informed consent from all study participants.

### Study population and ART procedures

In total 255 couples undergoing ART were included in the study from April 2018 to May 2019 from four fertility clinics in Sweden: IVF Stockholm, and the reproductive medicine units at Örebro University Hospital, Uppsala University Hospital and Karolinska University Hospital. All couples undergoing In vitro fertilisation (IVF) or intracytoplasmic sperm injection (ICSI) with single embryo transfers regardless of indication for the treatment were eligible to participate in the study. IVF or ICSI treatments utilising donor sperm or donor oocytes were excluded due to ethical concerns, as study specific informed consent could not be obtained from the donor.

Controlled ovarian stimulation was performed using either a short gonadotropin releasing hormone (GnRH) antagonist protocol or long GnRH agonist protocol. Gonadotropin administration involved recombinant follicle stimulating hormone or human menopausal gonadotropin, followed by oocyte maturation triggered with either recombinant hCG or, in selected cases, a GnRH agonist. Aspirated oocytes were fertilised in-vitro using either standard IVF or ICSI, according to the clinical routines and protocols used at each individual fertility clinic. Embryos were cultured in individual drops of either G-TL (Vitrolife, Sweden) or SAGE (Cooper Surgical, Denmark) single-step culture media at 37°C with 6% CO_2_ and 6% O_2_.

### Morphological assessment of the embryos

Morphological quality assessment of the embryos has been performed according to the Istanbul ESHRE consensus of 2011^27^. Cleavage stage embryos have been evaluated based on the number of blastomeres and the degree of fragmentation while blastocysts were scored based on the Gardner system that combine the developmental stage (blastocoel expansion and hatching status) with the appearance of the inner cell mass (ICM) and trophectoderm (TE)^28^. The developmental stage was scored from 1 to 6, with 6 corresponding to a completely hatched blastocyst. The ICM and TE were graded from A to C, where A corresponds to the highest grade (top quality). To be considered for embryo transfer on day 3, cleavage stage embryos had to have 6-10 blastomeres and ≤ 20% fragmentation whereas blastocysts had to have a minimum Gardner score of 3BB.

### Collection of spent culture media and blastocoel fluid

ESM were collected from transfer-quality cleavage stage embryos on culture day 3 and from blastocyst embryos on culture day 5 or 6. Collection of ESM was performed immediately after fresh embryo transfer or vitrification. A volume of 15-20 µl of ESM was aspirated by carefully avoiding the overlaying oil and transferred to a sterile cryotube. BF was collected from blastocysts undergoing laser collapsing as a routine procedure before vitrification to avoid ice crystal formation. To collect BF, the embryo was transferred to a new drop of culture media where laser collapsing of the blastocoel cavity was performed resulting in a release of BF into the culture media. A volume of 15-20μl of the BF containing culture media was collected in a sterile cryotube immediately after the collapsed embryo had been removed and vitrified. Culture media not exposed to an embryo but subjected to the same culture conditions and culture time, were collected as negative (medium only) controls from each new batch of culture media used in the study. All samples were immediately stored in the vapour phase of liquid nitrogen until further processing.

### Primary and secondary clinical outcomes

The primary clinical outcome used in our study was embryo implantation. A successful implantation was defined as a positive urine hCG measured 14 days after embryo transfer. As secondary outcomes we evaluated live birth and negative pregnancy outcomes defined according to the guidelines published by the International Committee for Monitoring Assisted reproduction technologies (ICMART) in 2017^29^. A live birth was defined as a birth occurring after 22 weeks of gestation, where the newborn exhibits signs of life, regardless of whether the umbilical cord has been cut or if the placenta remains attached. The negative pregnancy outcome group included the following pregnancy outcomes: (1) biochemical pregnancy, defined as detection of hCG after embryo transfer, but without any visible gestational sac on ultrasound performed week 7 to 9 of gestation; (2) Miscarriage, defined as spontaneous loss of a previously ultrasound verified intrauterine pregnancy before 22 weeks of gestation. Ectopic pregnancies were not included among the secondary outcomes, but the information on the outcome was collected and is presented in Extended Table 1.

### RNA isolation

The cryotube containing ESM or BF was briefly centrifuged at 567 × g for 1 minute. The content of each sample was then transferred to an individual RNAse-free 1.5 ml Eppendorf tube (Sigma Aldrich, Sweden). Next, 100 μl of Guanidium-based RLT plus buffer (Qiagen, Germany) was mixed with the sample followed by addition of 2 μl of pellet paint to visualise the RNA pellet. To precipitate the RNA, 160 μl of absolute ethanol was subsequently added to the sample followed by vortexing and incubation for 48 hours at −20°C. After incubation, the tubes were centrifuged at 16000 × g at 4°C for 20 min. The supernatant was then carefully discarded without disturbing the pink pellet. The RNA pellet was washed two times with 1ml of 80% fresh ethanol, air dried for 5 minutes and finally re-suspended in 7 μl of RNase free water and stored at −80°C until further processing.

### Polyadenylated RNA library preparation and sequencing

To detect cell-free polyadenylated RNA molecules in ESM and BF samples, we modified the Smart-seq3 cDNA preparation protocol for low-input biofluids ^30^. Briefly, 3.8μl of extracted total RNA was added as input volume per sample to a 0.5μl lysis buffer mix resulting in total volume of 4.3μl with final concentrations of 0.07% of Triton-X-100, 0.37 unit/μl of Recombinant RNase Inhibitor (Takara Bio, Japan), 0.87mM each of dNTP, and 1.4μM of Oligo dT. The mix was incubated at 70°C for 3 minutes. Water was included as negative controls. Reverse transcription was followed by adding 1.7μl of the reverse transcription mix resulting in total volume of 6μl with final concentrations of 5 unit/μl of Maxima H Minus reverse transcriptase enzyme (Thermo Scientific), 0.8 unit/μl of Recombinant RNase Inhibitor (Takara Bio, Japan), 50 mM of Tris-HCl pH 8.3, 75 mM of NaCl, 1.25 mM of GTP, 2.5mM of MgCl2, 5 mM of dTT, and 2μM of template-switching oligo. The reaction was incubated at 37°C for 30 minutes followed by 42°C for 1 hour, and 10 cycles of 2 min at 42°C and 2 min at 50°C and ended at 85 °C for 5 minutes. Post reverse transcription, PCR amplification was performed using 10μl of PCR mix resulting in total volume of μl with final concentrations of 1X of KAPA HiFi HotStart ReadyMix (Roche), 0.8μM of forward primer, and 0.1μM of reverse primer. The mix was incubated at 98°C for 3 minutes, followed by 20 cycles of 98°C for 20 seconds, 65°C for 30 seconds, and 72°C for 6 minutes, and ended at 72 °C for 5 minutes. The amplified libraries were purified with Agencourt AMPure XP beads (Beckmann Coulter, USA) in a 0.7 to 1 ratio of beads to sample and finally eluted in 10μl of RNAse free water after purification. The purified libraries were quality checked using a High Sensitivity DNA Kit (Agilent technologies, USA) on an Agilent 2100 Bioanalyzer instrument (Agilent technologies, USA). The cDNA quantity was measured with a Qubit flex Fluorometer (Invitrogen, Singapore) using the Qubit 1X dsDNA High Sensitivity Kit (Invitrogen, Oregon, USA) as per the manufacturers protocol. Tagmentation of the amplified libraries was performed using Illumina Nextera XT kit (Illumina Inc, USA) as per the manufacturers recommendations with custom made Nextera XT dual index primers (IDT technologies, Germany). After tagmentation, each sample was purified with AMPure XP beads in a 1:1 ratio of beads to sample followed by quality and quantity evaluation using a bioanalyzer and Qubit flex Fluorometer respectively using the same reagents as described above. Finally, 5ng of cDNA from each sample was pooled and sequenced on an Illumina Novoseq™ 6000 SP flow cell with a 100 base-pair single-end read setup.

### Bioinformatic analyses

#### Processing of RNA sequencing data

In total we performed RNA sequencing on 435 ESM and 148 BF samples. We filtered out 91 ESM and 72 BF samples from embryo that had not been transferred at the time of the analyses or were transferred on culture day 4. The raw FASTQ files were quality-checked using FastQC (Version 0.11.9). Read alignment, annotation and quantification were performed according to the previously published zUMIs pipeline (version 2.9.7) that was developed to handle UMI and sample specific barcodes and to allow for down sampling of hugely varying library sizes^31^. The pipeline was run using the suggested miniconda environment. The demultiplexed reads generated for each sample were merged using the script ‘merge_demultiplexed_fastq.R’. The merged fastq file was subjected to the zUMI pipeline. The samples were processed using a pre-defined YAML configuration file against human reference GRCh38 and Gencode v44, with the following additional modifications to STAR aligner ‘--clip3pAdapterSeq TGTCTCTTATACACATCT --outFilterMismatchNmax 5 --outFilterMatchNmin 5’. In brief, the reads were filtered with a Phred score cut-off of >20 to filter out low quality reads, resulting in removal of 9 ESM and 4 BF samples. The remaining reads were mapped to the human genome 38 (hg 38) using STAR aligner (version 2.7.3a), followed by counting and UMI deduplication. The gene counts were then stored in an RDS file. Counts from exonic reads with UMIs were selected from the RDS file for downstream analysis. Tin.py module from RSeQC was used to generate TIN values which was used to evaluate the RNA integrity of each sample^17^. An RNA molecule was considered as detected if at least 5 unique reads aligned to the gene. The final set of samples used for down-stream analyses consisted of ESM from 105 cleavage stage and 230 blastocyst stage embryos that had undergone fresh or frozen transfers, and BF from 72 blastocyst embryos that had underwent frozen transfers.

#### Differential gene expression

To identify differentially abundant polyadenylated RNAs in our data, we used the Bioconductor software DESeq2 (version 1.44.0). Differentially abundant genes (DAGs) with a p-value <0.05 and a log_2_ fold change of ≤ −1 or ≥ 1 were considered statistically significant. The DESeq2 analyses was performed between the following two groups: (1) hCG negative and positive embryos and (2) negative pregnancy outcome and live-birth embryos. ESM and BF samples were analysed separately. For the ESM samples, cleavage and blastocyst embryos were analysed separately. Further sub-analyses were also performed for blastocyst embryos according to developmental stage and culture day of transfer or vitrification to understand their influence on the results. Sub-grouping of the BF samples according to the day of transfer or vitrification and developmental stage deemed not feasible due to the small sample size. As a subgroup analyses, BF samples were stratified by maternal age, with samples from women older than 34 years analysed separately from those of women aged 34 years or younger.

#### Machine learning

The 72 BF samples were divided into one training and testing set, and one independent validation set. We randomly selected 22 samples for the validation set. To ensure a balanced representation of hCG-positive and hCG-negative cases, 11 hCG-positive and 11 hCG-negative samples were randomly selected from their respective groups. The remaining 50 samples were used for training and testing. The gene expression matrix was filtered to only include genes with counts of at least 1 across all samples.

Leave-one-out cross validation (LOOCV) was performed on the training and testing set to identify an optimal prediction model over all possible permutations of the following three main parameters of interest: (1) model class, (2) pre-processing, and (3) feature set. The model classes tested were random forest (RF), L2 Penalised Logistic Regression (L2-PLR) using stepPlr (version 0.93), and L1 Regularised Logistic Regression (L1RLR) using Glmnet (Matrix, version 1.7-0)^32–34^. Pre-processing was performed using scaling or principal component analysis (PCA). The feature set used as input for the model was either the complete gene set, or a filtered gene set based on DAGs. Morphological grading and maternal age were then added to the gene set as additional features in the model. Raw counts were used as input for the genes. Only genes detected in > 6 samples with minimum of 5 counts were included. DAGs were identified using DESeq2 (version 1.44.0) with a p-value of 0.1 as cut-off for significance.

The performance of the prediction models was evaluated using ROC curves and calculation of AUC, accuracy, specificity and sensitivity based on the confusion matrix generated. All analyses were performed using R version 4.4. The final selected parameters for the model were: L1RLR as model class, scaling as pre-processing of the count data and an input feature set that included raw counts from all genes and maternal age. The abundancy of the genes contributing with weightage in the model were checked in the control media. Genes with > 5 counts in any of the control samples were filtered out, resulting in exclusion of three genes from the model.

## Acknowledgements

We want to acknowledge all the patients that have contributed with invaluable material for this study and all the midwives, physicians and embryologists that have kindly assisted with the sample collection at each fertility clinic. In addition, we want to acknowledge the methodological support provided by SciLifeLab’s Bioinformatics platform (NBIS) and by Clinicum at Karolinska University Hospital.

## Author contributions

ORF, KGD and PGL supervised the study. KGD was responsible for overall study conduct, regulatory approvals, conception, and funding. NRB contributed with the original research idea and conception. PGL contributed with the original research idea, with insights on clinical embryology, conceptualised and led the study. PGL and ORF did the feasibility studies. KGD, NRB, PGL, CF and ORF designed the study. CF, KGD and PGL obtained the regulatory permits. KGD, PGL, NRB, ORF, CF and JM developed the sample collection protocol and planned the sample collection. CF and NRB coordinated the multicentre sample collection. WA, MS, KL, VT, JIO, JM and TB were responsible on-site for the sample collection at each individual fertility clinic. NRB and ORF developed the RNA extraction protocol. CF, NRB and ORF developed the RNA library preparation protocol for ESM and BF samples. NRB, ORF and CF designed the final sequencing experiments and NRB constructed the cDNA libraries. CF performed cleaning and quality controls of the clinical data. KN performed the bioinformatic analyses of the RNA sequencing data. FK created the machine learning model with supervision from FV and KN refined it. CF, PGL, ORF and NRB designed the data analysis, provided continuous scientific input and guidance during the data analyses. CF, KN, NRB, ORF, KGD and PGL interpreted the results. CF and ORF drafted the manuscript. All authors critically reviewed, edited and approved on the final version of the manuscript.

---

**Extended Fig. 1.**
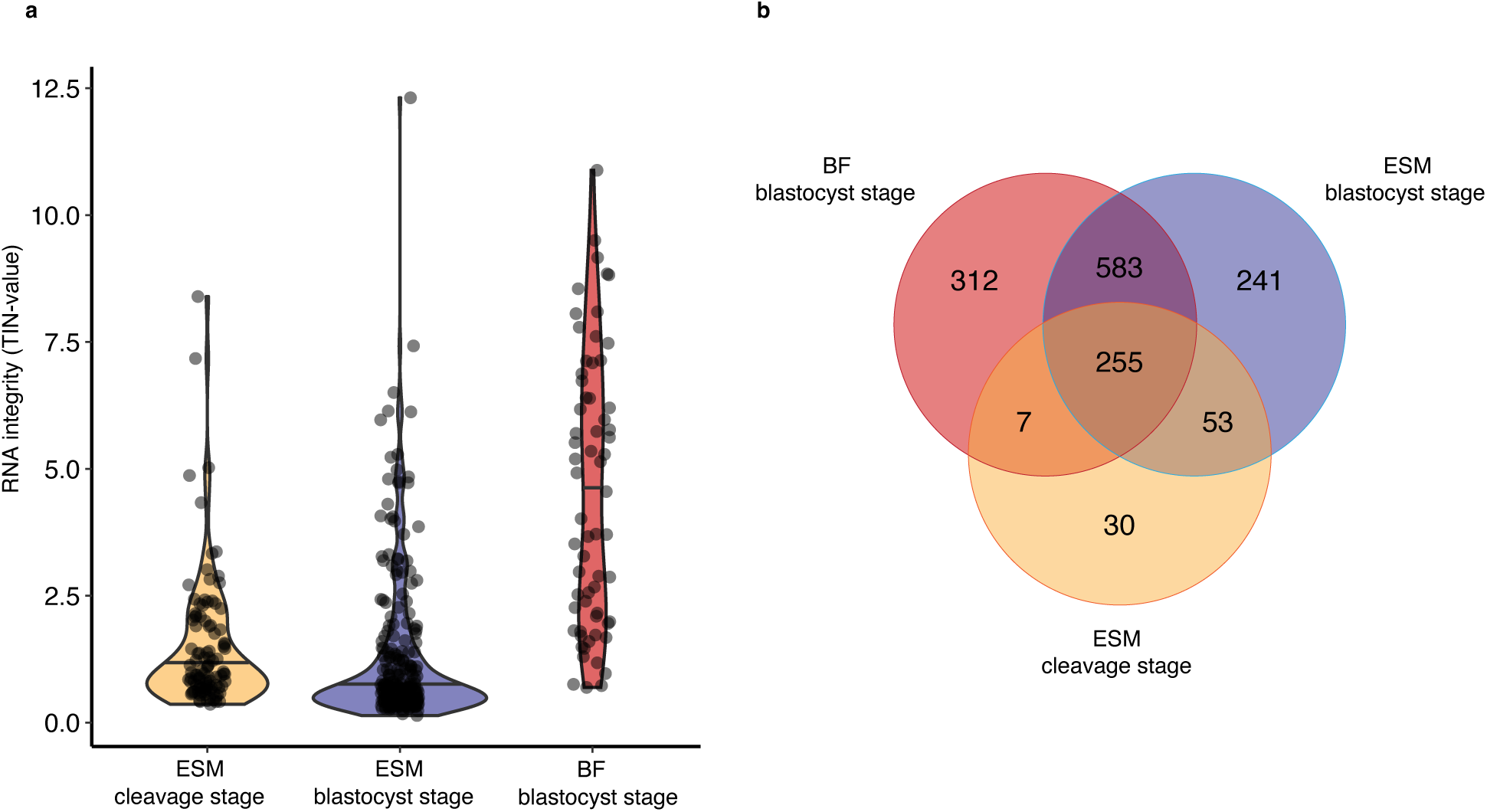
RNA integrity and overlap of detected genes in ESM and BF. **a**, Violin plot showing the RNA integrity as TIN values for BF samples from transferred blastocyst stage embryos (n=72) and ESM samples from transferred blastocyst-(n=230) and cleavage-stage embryos (n=105). **b**, Ven diagram showing the overlap of detected genes in the ESM samples from cleavage- and blastocyst-stage embryos and in the BF samples from blastocyst stage embryos. Abbreviations: TIN=transcript integrity number; ESM=embryo spent media; BF=blastocoel fluid.

**Extended Fig. 2.**
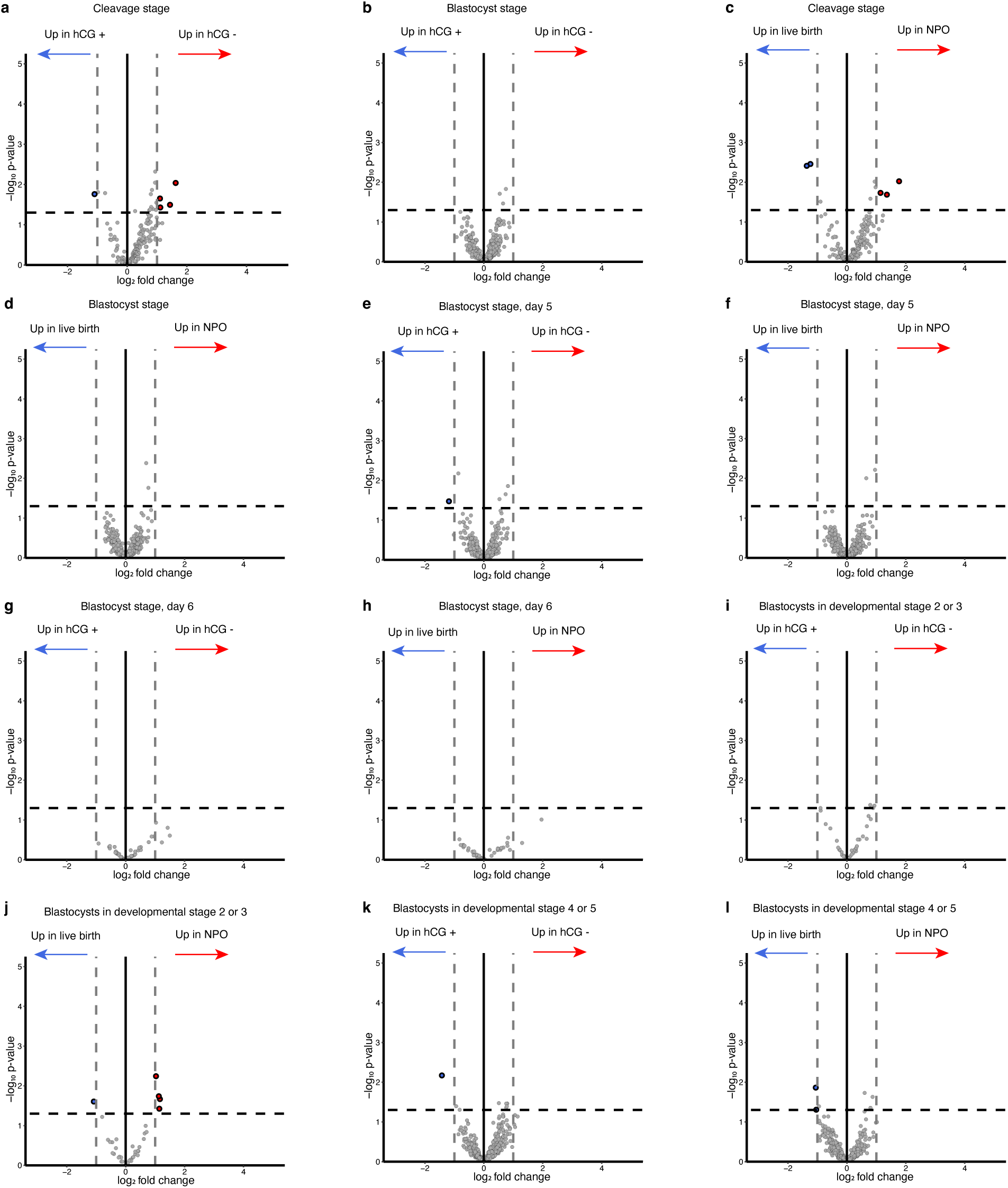
Detection of polyadenylated RNA associated with pregnancy outcomes in embryo spent media. **a-l**, Volcano plots demonstrating differences in RNA levels in embryo spent media from embryos transferred at cleavage stage (**a & c**) and at blastocyst stage (**b & d-l**) when comparing hCG negative and hCG positive outcomes as well as NPO and live births. The analysis of blastocyst embryos has included all blastocyst embryos (**b & d**), or sub-groups of blastocyst embryos based on embryo culture day at transfer or vitrification (**e-h**) and the developmental stage of the blastocyst embryo according to morphological grade (**i-l**). Red and blue dots represent significantly upregulated and downregulated genes respectively detected using DESeq2 (p-value <0.05 and a log2 fold change ≤-1 or ≥1). Grey dots are unsignificant genes. Abbreviations: ESM=embryo spent media; hCG=human

**Extended Fig. 3.**
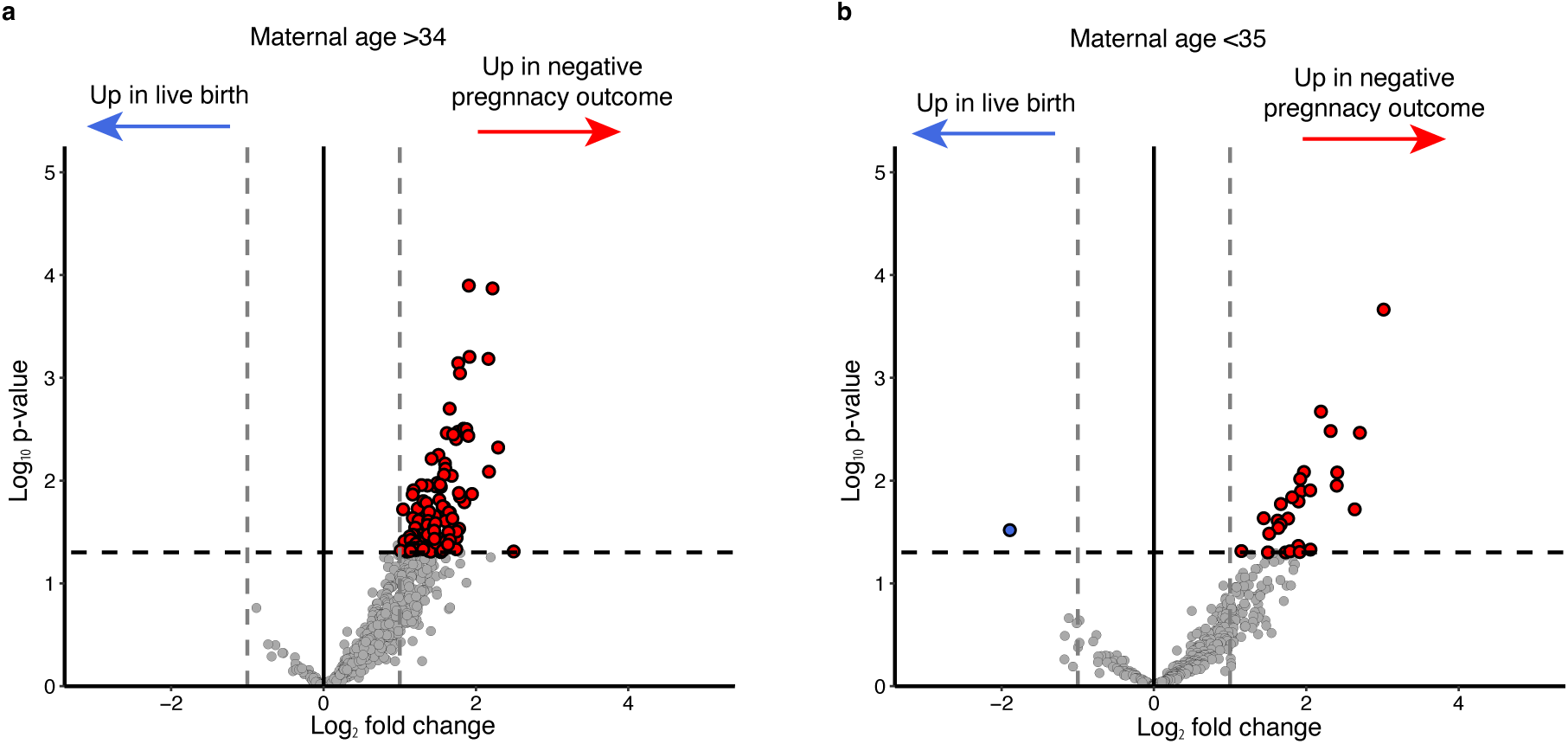
Detection of pregnancy outcome associated RNA in blastocoel fluid stratified by maternal age. **a-b**, Volcano plots demonstrating differences in blastocoel fluid RNA between negative pregnancy outcome and live birth blastocysts from women old than 34 years (n=27) (**a**) and younger than 35 years (n=41) (**b**). Red and blue dots represent significantly upregulated and downregulated genes respectively detected using DESeq2 (p-value <0.05 and a log2 fold change ≤-1 or ≥1). Abbreviations: hCG=human chorionic gonadotropin

**Extended Fig. 4.**
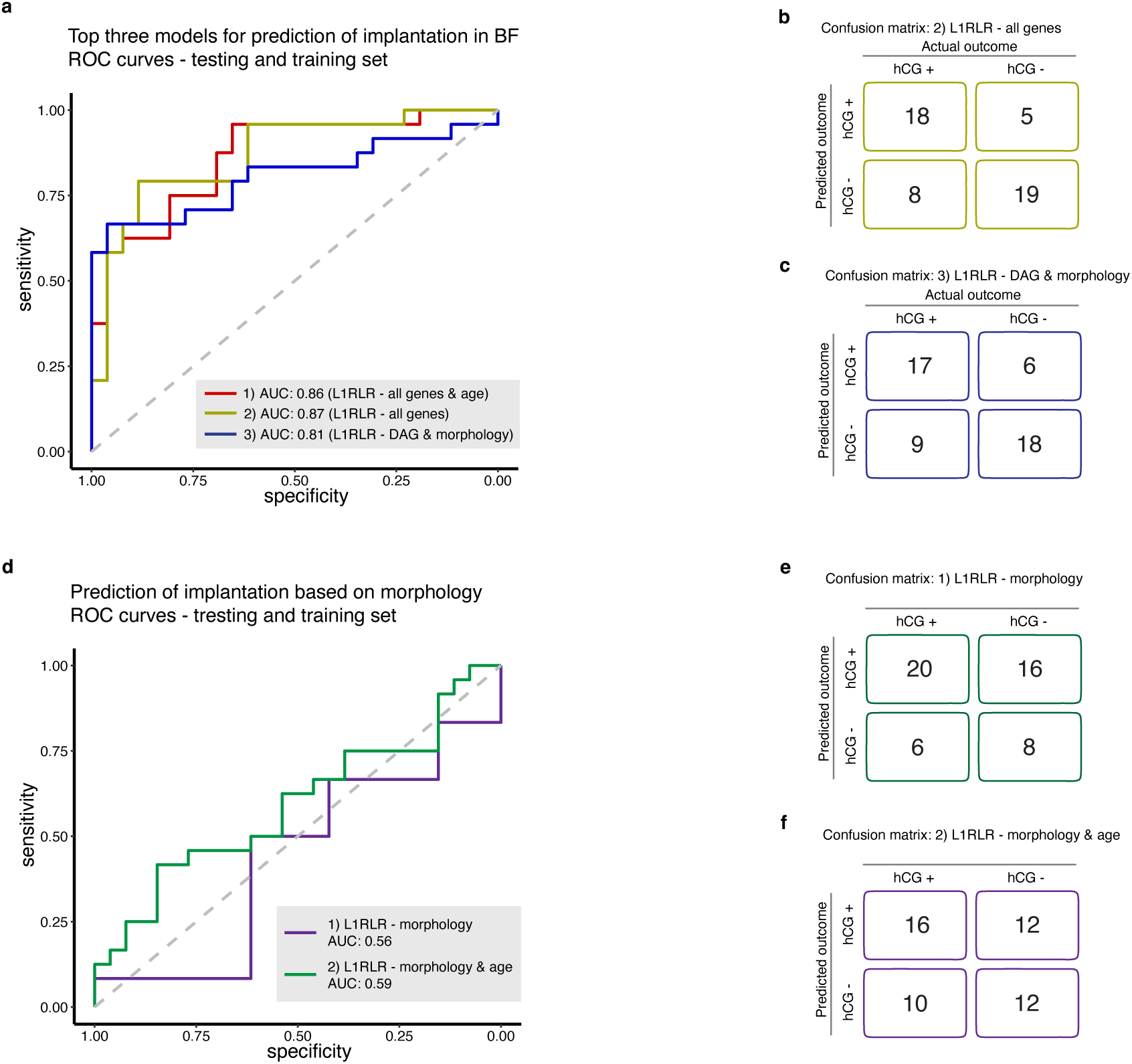
Evaluation of model performance with different input features. **a**, ROC curves of the three most accurate prediction models of implantation outcome based on RNA in BF in the training and testing set. The three prediction models utilise L1RLR as model class, scaling for pre-processing and the following differences in input features: (1) all genes and maternal age; (2) all genes, and (3) differentially abundant genes and morphological scores. Differentially abundant genes were identified using DESeq2 with a p-value <0.01 as cut-off for inclusion in the model. **b-c**, confusion matrices of the second (**b**) and third (**c**) most accurate prediction models. The confusion matrix of the most accurate model is shown in figure 2. d, ROC curves for the prediction models of implantation outcome based on morphological scores of the embryos included in the BF cohort. The two evaluated models incorporated L1RLR as model class with morphological scores alone (1) or combined with maternal age (2) as input features. **e-f**, Confusion matrix of the morphology alone (**e**) and morphology combined with maternal age (**f**) prediction models. L1RLR was performed with Glmnet. Abbreviations: BF=blastocoel fluid; hCG=human chorionic gonadotropin; L1RLR=L1 regularised logistic regression.

**Extended Table 1.** Clinical characteristics of the transferred embryos included in the data analysis.

| Parameter | ESM<br>Cleavage stage |  | ESM<br>Blastocyst stage |  | BF<br>Blastocyst stage |  |
| --- | --- | --- | --- | --- | --- | --- |
|  | hCG<br>positive<br>(n= 42) | hCG<br>negative<br>(n= 63) | hCG*<br>positive<br>(n= 111) | hCG<br>negative<br>(n=119) | hCG*<br>positive<br>(n=37) | hCG<br>negative<br>(n=35) |
| Maternal age (mean) | 34 | 34 | 32 | 33 | 32 | 34 |
| Fresh embryo transfer (%) | 93% | 90% | 42% | 49% | 0% | 0% |
| Frozen embryo transfer (%) | 7% | 10% | 58% | 51% | 100% | 100% |
| IVF (%) | 62% | 67% | 66% | 63% | 70% | 66% |
| ICSI (%) | 38% | 33% | 34% | 37% | 30% | 34% |
| Number of IVF cycle attempts (mean) | 2 | 2 | 2 | 2 | 2 | 2 |
| Culture day 5 at transfer /freezing (%) | --- | --- | 92% | 87% | 86% | 83% |
| Culture day 6 at transfer /freezing (%) | --- | --- | 8% | 13% | 14% | 17% |
| Biochemical pregnancy (%) | 2% | --- | 7% | --- | 0% | --- |
| Miscarriage (%) | 14% | --- | 16% | --- | 16% | --- |
| Ectopic pregnancy (%) | 5% | --- | 0% | --- | 0% | --- |
| Live birth (%) | 79% | --- | 72% | --- | 73% | --- |

\*Data on pregnancy outcome is missing in five and four ESM and BF samples from hCG positive blastocyst stage embryos respectively. Abbreviations: ESM=embryo spent media; BF=blastocoel fluid; IVF=in-vitro fertilization; ICSI=intracytoplasmic sperm injection; hCG=human chorionic gonadotropin.

**Extended Table 2.**
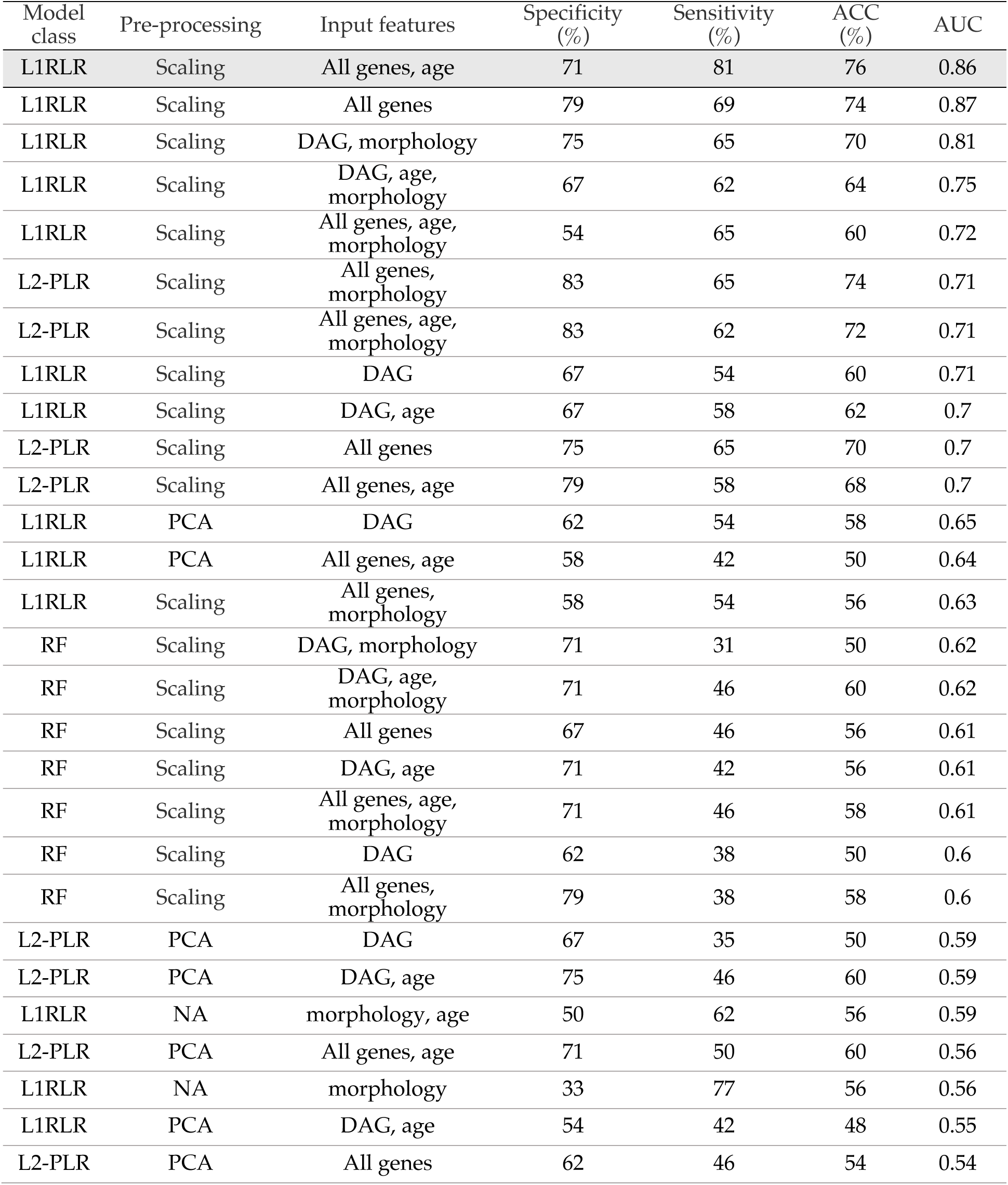
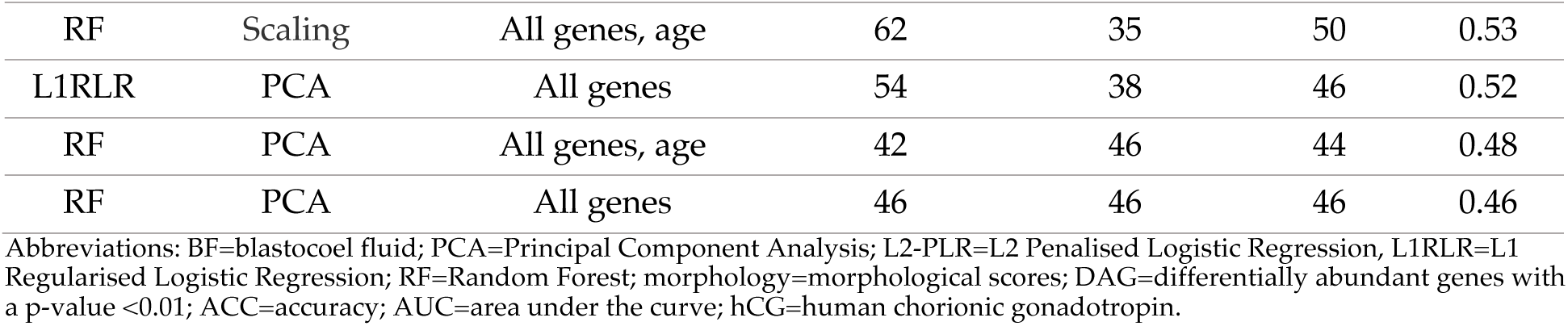
Evaluation of models to predict implantation outcome after transfer of blastocyst stage embryos. The results are based on the training and testing set. Model classes evaluated included RF, L1RLR using Glmnet, and L2-PLR using stepPLR. Pre-processing was performed with PCA or scaling. Polyadenylated RNA in BF, maternal age, and morphological scores of the embryos were evaluated as input features in different combinations. The input feature “all genes” means that no filtering using differential expression analysis was performed on the BF RNA. Implantation was defined as a positive urine hCG 14 days after embryo transfer.

